# Antimicrobial exposure in sexual networks drives divergent evolution in modern gonococci

**DOI:** 10.1101/334847

**Authors:** Leonor Sánchez-Busó, Daniel Golparian, Jukka Corander, Yonatan H. Grad, Makoto Ohnishi, Rebecca Flemming, Julian Parkhill, Stephen D. Bentley, Magnus Unemo, Simon R. Harris

**Author notes:** Corresponding author: Simon R. Harris.

## Abstract

The sexually transmitted pathogen *Neisseria gonorrhoeae* is regarded as being on the way to becoming an untreatable superbug. Despite its clinical importance, little is known about its emergence and evolution, and how this corresponds with the introduction of antimicrobials. We present a genome-based phylogeographic analysis of 419 gonococcal isolates from across the globe. Results indicate that modern gonococci originated in Europe or Africa as late as the 16^th^century and subsequently disseminated globally. We provide evidence that the modern gonococcal population has been shaped by antimicrobial treatment of sexually transmitted and other infections, leading to the emergence of two major lineages with different evolutionary strategies. The well-described multi-resistant lineage is associated with high rates of homologous recombination and infection in high-risk sexual networks where antimicrobial treatment is frequent. A second, multi-susceptible lineage associated with heterosexual networks, where asymptomatic infection is more common, was also identified, with potential implications for infection control.

---

Almost 360 million curable sexually transmitted infections (STIs) are estimated to occur globally each year, with *Neisseria gonorrhoeae*, the causative agent of gonorrhoea, infecting approximately 78 million^1^. The highest gonorrhoea burden is among men, although problematic infections are more common in women for whom the early stages of infection are often asymptomatic. Unresolved urogenital infections can lead to severe complications and sequelae, such as reproductive problems including infertility, serious eye infections in newborns, and enhanced transmission of HIV^2^. The emergence and proliferation of gonococci with reduced susceptibility to front-line antimicrobials such as extended-spectrum cephalosporins (ESCs; cefixime and ceftriaxone) and azithromycin have contributed to, although do not completely explain, the increase in incidence of gonorrhoea. Resistance to dual therapy (injectable ceftriaxone plus oral azithromycin), the current recommended treatment in many countries, is fortunately rare^3^, however, ceftriaxone resistance has been reported from all continents and azithromycin resistance is on the increase globally^4^, raising fears that the effectiveness of this regimen will be short-lived. Much of the focus of gonococcal control is on particular high-risk sexual networks that often partake in unprotected sex with multiple partners, particularly sex workers and men who have sex with men (MSM) but also young heterosexuals. These groups are more frequently exposed to both infection and antimicrobial treatment, which has led to these networks being the suspected drivers of antimicrobial resistance (AMR)^5^. However, AMR is not the only factor driving the recent success of *N. gonorrhoeae*. Dual therapy is effective against the vast majority of infections, yet since its introduction, gonorrhoea infections have continued to increase in most settings^6^.

Georges Luys famously opened his medical textbook on gonorrhoea with the statement that ‘Gonorrhoea is as old as mankind’^7^. However, despite *N. gonorrhoeae* often being described as an ancient pathogen, there are no clear descriptions of a disease like modern gonorrhoea in the ancient sources. Some compatible symptoms do appear in the medical literature of classical Greece and Rome, but nothing decisive, and the presence or absence of modern gonorrhoea in the ancient Mediterranean has been much debated as a result^8^. Early modern terms like ‘the clap’, ‘the pox’, or ‘the venereal disease’ also covered a range of conditions, and it was not until 1879 that Alfred Neisser identified the bacteria that now bears his name^9^. AMR in gonorrhoea became apparent soon after antimicrobials were first introduced for its treatment. Sulfonamide therapy was initially successful, but shortly after its introduction in 1935 the emergence of resistance curtailed its effectiveness. Subsequently, gonococci have developed resistance to all available treatments, including penicillins, tetracyclines, quinolones, macrolides and ESCs^2^. One characteristic of *N. gonorrhoeae* that has played an important role in its rapid gain and spread of AMR is its ability to exchange DNA via homologous recombination both within its own species and with another *Neisseria*. For example, mosaic penicillin-binding protein 2 (PBP2; encoded by the *penA* gene) alleles gained via recombination have been key in the emergence of resistance to ESCs^10,11^ which led to the replacement of cefixime as the first-line treatment for gonorrhoea. The first ceftriaxone resistant mosaic *penA* genotype was seen in an isolate from a pharyngeal infection in a female sex worker in Japan in 2009^12^, but similar mosaics were subsequently soon seen worldwide^2,13^. In fact, a number of resistances have first been identified in Japan, leading to the hypothesis that most AMR gonorrhoea originates there, or elsewhere in the WHO Western Pacific Region^2^.

Whole-genome sequencing has been successfully used to reveal the origins, global spread and population structure of several human pathogens^14^. However, gonococcal genome sequencing has mostly targeted specific populations and outbreaks^15^-^19^. Here, we report the findings of a global genomic study of 419 *N. gonorrhoeae* isolates spanning five continents and more than 50 years, including varying susceptibilities to important antimicrobials. Our aim was to elucidate when and where modern gonococcal populations emerged, evolved and dispersed, and how antimicrobial usage and transmission in different sexual networks has influenced their population dynamics.

## Modern gonococcus not ‘as old as mankind’

Our collection spans a period of more than 50 years (1960 – 2013) and 58 countries from five continents (Supplementary Table 1, Figure 1). A population-level analysis revealed a high level of admixture among *N. gonorrhoeae* with no significant differentiation between continents (Supplementary Table 2), with the exception of Africa (Extended Data Figure 1, Supplementary Table 3-5). We estimated the substitution rate for the non-recombining section of the genomes in the collection (Extended Data Figure 2, panel a) to be 3.74E^-06^ substitutions/site/year CI (confidence interval) [3.39E^-06^ – 4.07E^-06^], which is similar to previous reports^15,17^ and comparable to rates calculated for other bacteria^20^. The time of the most recent common ancestor (tMRCA) was estimated to be around the 16^th^ century (1589, CI [1544 – 1623]) (Figure 2). Although high rates of recombination can lead to underestimation of tMRCAs to some extent, these results are strongly at odds with the hypothesis that modern gonorrhoea has existed as long as mankind and cast further doubt on the ascribing of historical descriptions of gonorrhoea-like symptoms to infection with ‘modern’ gonococci.

**Figure 1.**
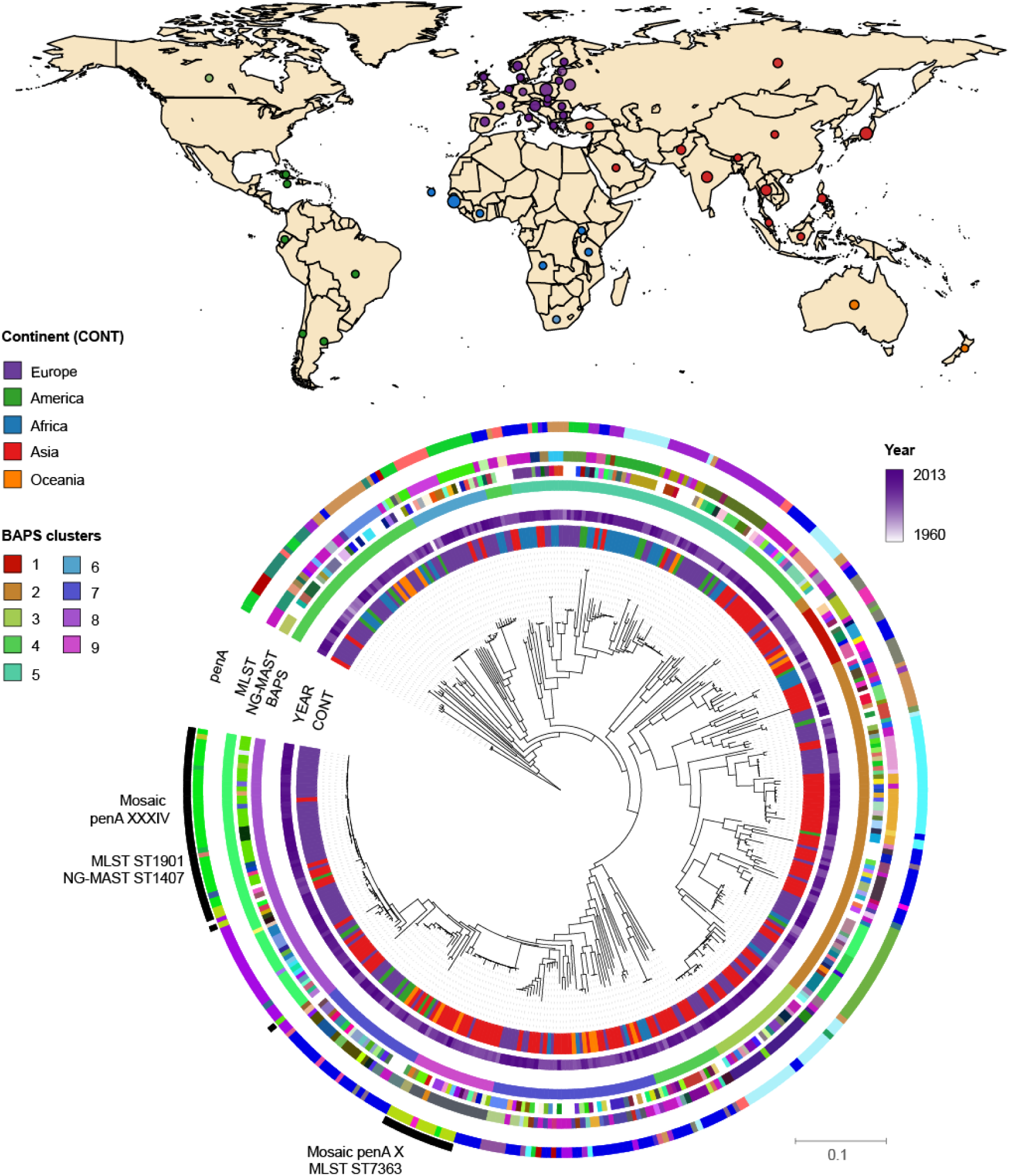
Geographic and phylogenetic distribution of *Neisseria gonorrhoeae* isolates. The map shows the countries of isolation of the strains in the collection coloured by continent. The phylogeny shows the relationship among the strains. Coloured strips show (from inside out) the continent of isolation (CONT), year and further typing information (BAPS clusters, NG-MAST, MLST and *penA* types; colours represent different types or alleles). Mosaic *penA* types are marked in the outermost black strip.

**Figure 2.**
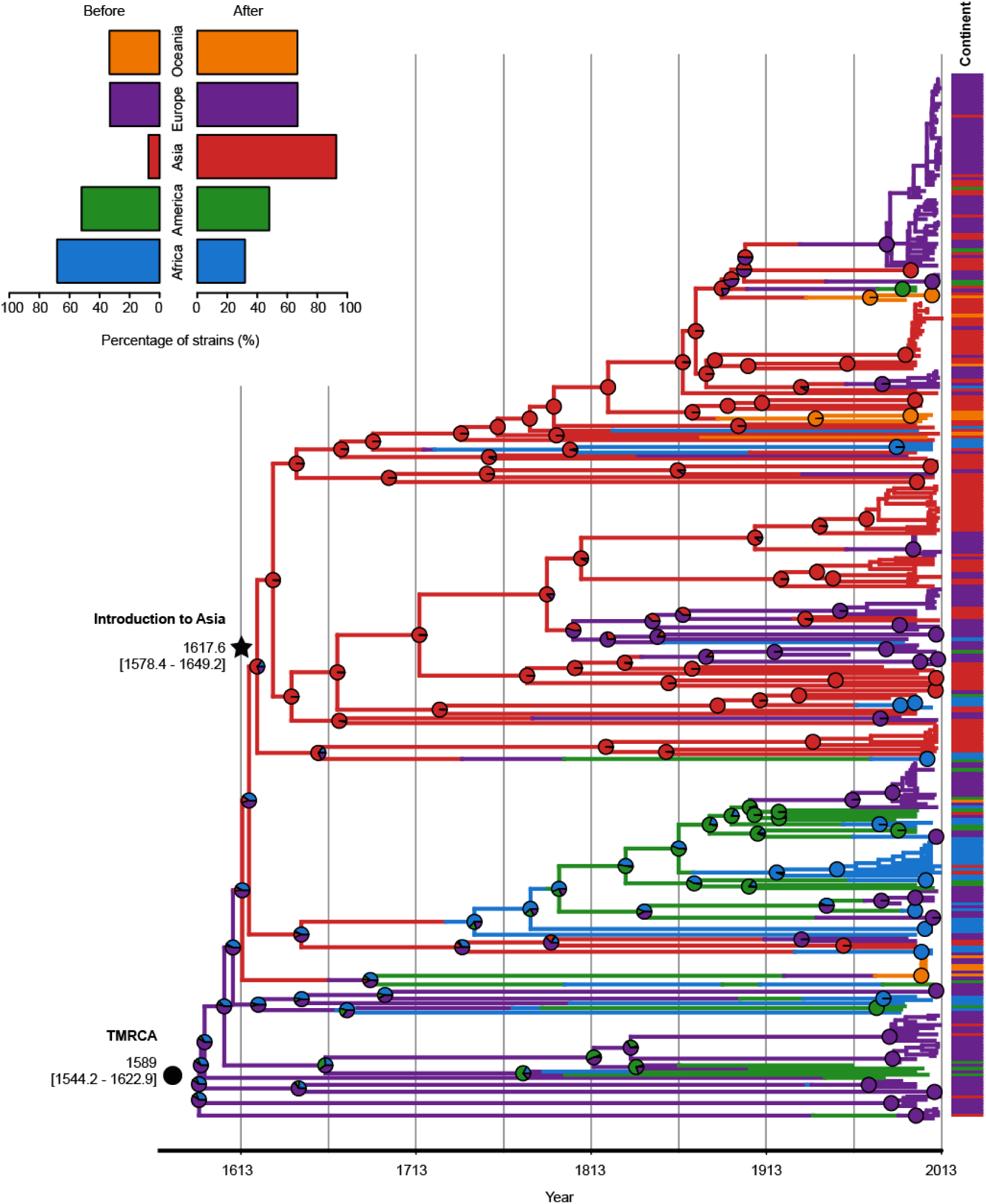
Global phylogeographic analysis. The dated maximum likelihood phylogenetic tree shows the posterior probabilities for each continent in every node (pie charts). Continents of isolation (prior) are shown as metadata next to the tips. The top left legend contains information on the proportion of strains from different continents before and after the introduction to Asia.

Despite modern gonococci being globally mixed, we found strong evidence of historic geographical separation, suggesting rapid mixing of populations is a relatively recent phenomenon. A phylogeographic analysis ascribed the origin of our collection to Europe (60.9% proportion of inferred ancestry). However, when corrected for biases in the number of samples from each continent, in part by complementing using isolates from a US study^15^, there was support for an African (90.7% proportion of inferred ancestry) origin (Extended Data Figure 3, panel a and b and Supplementary Table 6). From this African root, we identified a number of change-points in the continental distribution of isolates across the tree (Extended Data Figure 3, panels c-d). The most significant change-point separated a basal lineage containing a high proportion (68.2%, 30/44) of African isolates and a lineage containing a high proportion of Asian isolates (92.6%, 137/148), despite the temporal sampling from the two continents being similar (Extended Data Figure 3, panels e-f). When combined with the dating, this can be interpreted as an early introduction of the modern gonococcus population into Asia (1617, CI [1578 – 1649], Figure 2) soon after its emergence in Africa or Europe. More recently, many re-introductions into the rest of the world have occurred from this Asian lineage, contributing to the highly mixed population observed today.

## Emergence of antimicrobial resistant gonorrhoea

Interestingly, MICs for six antimicrobials (Extended Data Figure 4) and the occurrence of genetic AMR determinants were significantly higher among the isolates belonging to the lineage that arose after the phylogeographic breakpoint representing the initial introduction into Asia (Student’s t p-value < 0.0001) (Figures 3 and 4c). We will therefore refer to the 298 after the breakpoint as lineage A and the 121 isolates before the breakpoint as lineage B.

**Figure 3.**
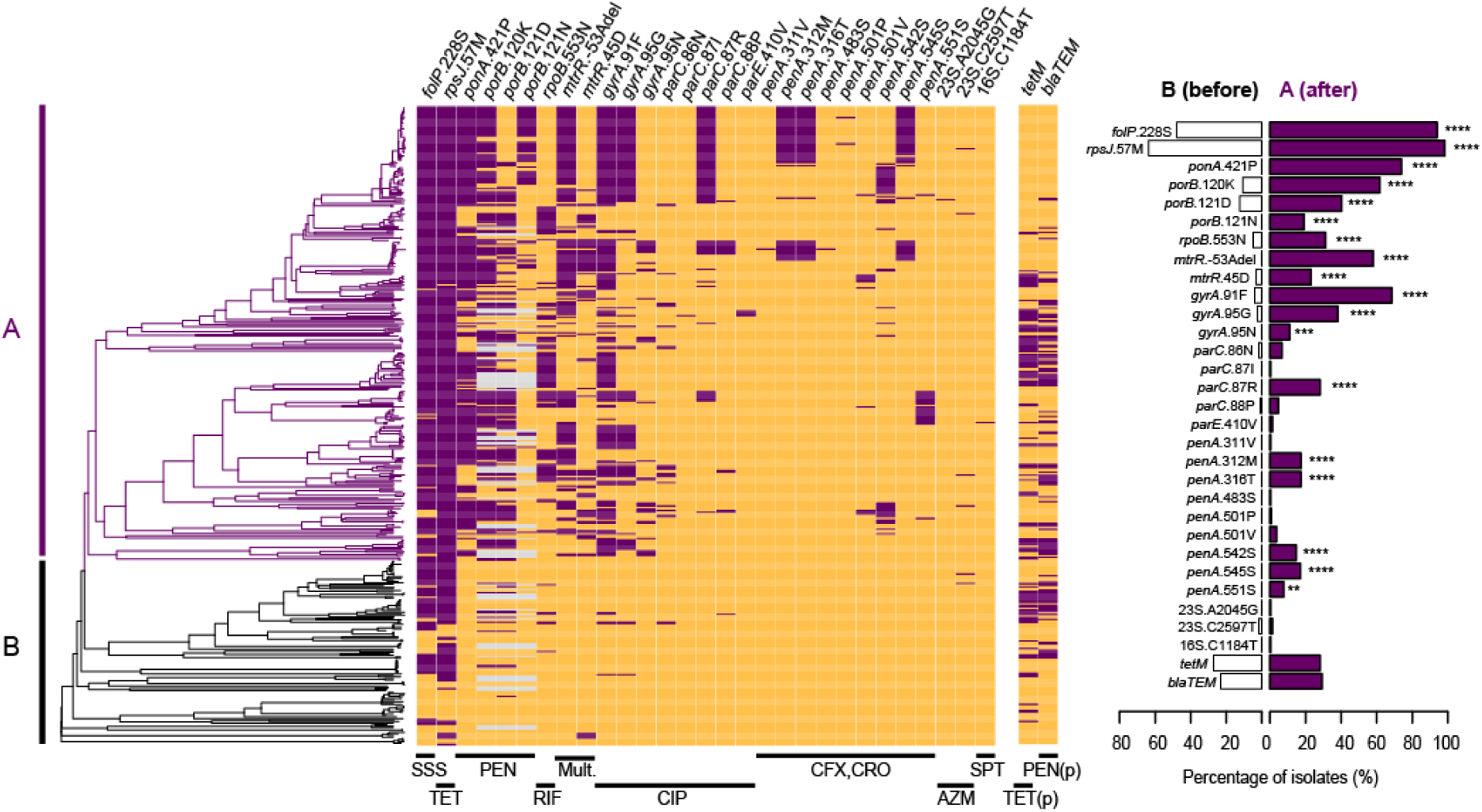
Evolution of antimicrobial resistance genetic determinants in *Neisseria gonorrhoeae*. Antimicrobial resistance determinants (chromosomal mutations and presence/absence of the *tetM* and *blaTEM* genes on plasmids (p)) detected in the whole dataset on the maximum likelihood dated tree. Purple represents presence of the determinant and orange its absence. Grey indicates isolates possessing *porB1a* rather than *porB1b*. The two main lineages are marked as A and B. The left graph shows the proportion of strains with each resistance determinant for both lineages. Statistical significance from a t-test is also shown in the graph. ****p-value<0.0001, ***p-value<0.001, **p-value<0.01, *p-value<0.05.

**Figure 4.**
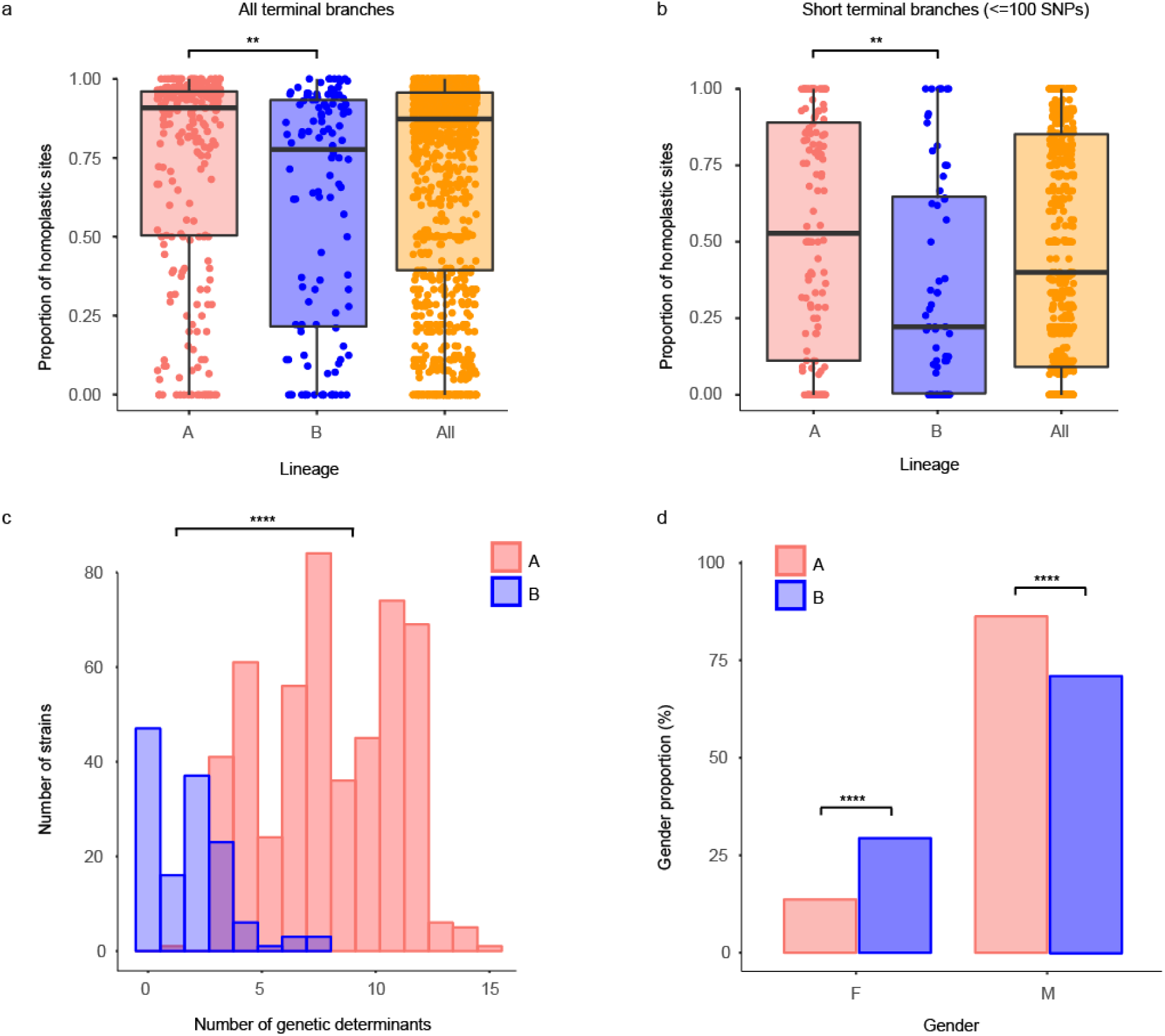
Characterization of the lineages of *Neisseria gonorrhoeae*. Proportion of homoplasic sites in all terminal (A) and short terminal branches (<=100 SNPs) (B) in lineages A and B and all strains. (C) Distribution of the total number of antimicrobial genetic determinants per strains in each lineage. (D) Proportion of strains isolated from female (F) and male (M) patients in each lineage. ****p-value<0.0001, **p-value<0.01.

Two AMR determinants, *folP* R228S, which reduces susceptibility to sulfonamides, and *rpsJ* V57M, which reduces susceptibility to tetracyclines, were carried by a large proportion of isolates, especially in lineage A (Supplementary Table 1, Figure 3). Fifty-one isolates contained a mosaic *penA* allele^21^, which are generally believed to be the result of recombination events with other *Neisseria* species. We identified three independent gains of mosaic alleles, all in lineage A. In a clade of 59 isolates with MLST ST1901, a first recombination event replaced the wild type allele with a *penA*10 allele and a subsequent event replaced *penA*10 with *penA*34. These two alleles differ by 16 SNPs and a codon insertion in the last 105 bases of the nucleotide sequence. Two isolates in this clade exhibited high Minimum Inhibitory Concentrations (MICs) for both cefixime (3-4 μg/ml) and ceftriaxone (2 μg/ml) (Extended Data Figure 5), and these were found to possess a *penA*42, which is a single SNP (A501P) variant from *penA*34^22,23^. In another lineage, associated with MLST ST7363, most isolates possessed the *penA*10 alleles, but we again observed a case of replacement with *penA*34. Only one isolate carried the A2045G 23S rRNA mutation (A2059G in *Escherichia coli* nomenclature) that confers high-level resistance to azithromycin. Six carried the low-level resistance C2597T 23S mutation (C2611T in *E. coli* nomenclature). As expected, isolates carrying a *tetM* gene on a conjugative plasmid and/or the *bla*^TEM^ plasmid showed increased resistance to tetracyclines and penicillin, respectively (Extended Data Figure 5). Strikingly, these two plasmids co-localized far more frequently than expected (Fisher’s test p-value < 0.0001), possibly reflecting the mobilization of pBlaTEM by the pConjugative plasmid^24^, and were completely absent from ESC-resistant isolates (Figure 3). The plasmid-encoded resistances showed no significant difference in prevalence in lineages A or B (*tetM* p-value = 0.92, *bla*^TEM^ p-value = 0.35). In contrast, of the 29 chromosomally-mediated resistance substitutions examined, 18 were significantly associated with clade A (Figure 3). Importantly, based on our phylogenetic dating, the majority of occurrences of these 29 determinants were estimated to have been acquired after the introduction of the antimicrobial against which they act (Extended Data Figure 6). The Gonococcal Genomic Island (GGI) was found in 277 (67%) isolates (Extended Data Figure 7), but showed no clear association with AMR.

## Two strategies for gonococcal success

Overall, our data show far fewer gains of chromosomally-encoded AMR determinants in lineage B compared to A (Extended Data Figure 8, panel a). Since these determinants primarily spread through the population via homologous recombination, such differences could be explained by differences in recombination frequency. To assess this, we compared the proportion of homoplasic sites, an indicator of recombination, in the terminal branches of the phylogenetic tree in the two lineages. This confirmed a significantly higher proportion in clade A, particularly for short branches, which represent very recent evolution (t-test p-value < 0.005; Figures 4a-b and Extended Data Figure 9). Similarly, the proportion of clustered SNPs, another signal of recombination, was also higher on the terminal branches in lineage A (p-value < 0.05).

One explanation for such differences could be opportunity. For recombination to occur, donor and recipient bacteria must co-localise. Thus, recombination between gonococci would be expected to occur more frequently in high-risk host populations where coinfection with other STIs and pharyngeal infections, which allow access to commensal *Neisseria* species, are more common. For gonorrhoea, these risk-groups include MSM, sex workers and some groups of young heterosexuals. These groups are also more likely to be exposed to repeated antimicrobial therapy for gonorrhoea and other STIs^5^. Unfortunately, due to limitations in availability of data on patient sexual behaviour, we could not adequately assess association of the lineages to risk factors in our dataset. However, we could analyse the distribution of the gender of the patients from which the isolates were taken. To increase the power of the analysis, we included 376 isolates from two North-American genomic studies^16,25^, to give a set of 639 isolates with complete gender information. Strikingly, lineage B included a significantly higher proportion of women (40/136, 29.4%) than A (69/503, 13.7%) (p-value < 0.0001) (Figure 4d and Extended Data Figure 8, panel b), which would suggest that B is more closely associated with heterosexuals. Corroborating this, data from a 2013 European-wide structured survey of over 1,000 *N. gonorrhoeae* isolates^26^ showed a similar pattern. Lineage B isolates were strongly associated with reduced MICs and female patients (61/214, 28.5% of lineage B isolates were from women vs 100/821, 12% of lineage A; p-value < 0.0001), and more importantly, of the patients that reported sexual orientation, 78.3% (94/120) of isolates in lineage B were from heterosexuals, in contrast to 52.6% (200/380) in A (p-value < 0.0001) (Extended Data Figure 10). Particular sublineages within lineage B appear particularly strongly associated with heterosexuals^26^. We suspect lineage A, being associated with higher-risk populations, does have greater opportunity for recombination, which may explain the observed higher recombination rate. However, transmission between low and high-risk populations is common within lineage A, so we suspect opportunity is not the only explanation for the differential recombination rate in the two lineages. The observation that plasmid-born resistances do not show the same difference in frequency between the two lineages also supports this view.

## Discussion

Gonorrhoea is one of the most clinically important STIs worldwide. Its rapid mode of transmission, especially among high-risk groups, and the emergence of resistance to many antimicrobials, has made the control of *N. gonorrhoeae* of primary importance for public health. In recent years there has been an understandable focus on AMR gonorrhoea, with resistance to all classes of antimicrobials used to treat the infection having been reported^2^. However, the increase in prevalence of gonorrhoea has continued in many settings^6^ despite resistance to dual therapy being extremely rare.

Our genomic analysis revealed a contemporary global population with little geographical structure, suggesting rapid recent intercontinental transmission is occurring. In particular, introductions from Asia into the rest of the world appear common, consistent with the observation that a number of recent resistant gonococcal clones have emerged from this region^2^. The one exception was Africa, where the sampled gonococcus was less diverse, potentially reflecting the relatively low rates of human migration with other continents^27^. However, our African sample size was small due to limited availability of isolates, so further study is required in this area.

We estimated an origin of modern gonococci in the 16^th^ century (1544 - 1623), which contrasts with historical interpretations of gonorrhoea as an ancient disease. Although we are keen to stress that high rates of recombination make accurate estimates difficult, and our confidence intervals are probably too narrow, this dating suggests that ancient accounts of gonorrhoeal-like symptoms may have been caused by other pathogens, or are evidence of an ancient *N. gonorrhoeae* population distinct from that observed today. It certainly disputes the view that the disease we now know as gonorrhoea is ‘as old as mankind’. The 16^th^ century was, nonetheless, an opportune time for the global dissemination of pathogens. It was a period of early modern globalization marked by the initiation and intensification of many intercontinental trade links, particularly by sea^28^. This period was of utmost importance for globalization due to an expeditious increase in exchange of goods, including the import of new crops from the Americas to Europe. Increased movement of people around the world also spawned local epidemics and pandemics^29^, and may well have played an important role in the evolution of modern gonorrhoea. A phylogeographic analysis using several subsampled set of strains from different continents to avoid bias placed the origin of the current global gonococcal population in Europe or Africa, and identified a subsequent introduction into Asia in the early 17^th^century (1578 – 1649), which expanded rapidly throughout the continent. More recently this lineage has been repeatedly transmitted back to the rest of the world.

A major finding is a strong association between isolates from the lineage that evolved from this early introduction to Asia and the development of AMR. Nearly all isolates in this lineage A, and over 50% of those lineage B, harboured resistance to sulfonamides (*folP* R228S mutation) and tetracyclines (*rpsJ* V57M mutation). Sulfonamides were the first antimicrobials introduced to treat gonorrhoea in 1935, with initial efficacies of around 90%. By the mid to late 1940s sulfonamide resistance was common, and it was discarded as a treatment for gonorrhoea^2^. However, sulfonamides are still widely used in combination with trimethoprim (TMP-SMZ) for prophylaxis in HIV positive patients and to treat a variety of bacterial infections^30^. Doxycycline (a tetracycline) is still sometimes used to treat gonococcal or presumptively non-gonococcal urethritis/cervicitis and is the recommended treatment for lymphogranuloma venereum^31^. We therefore suspect the high incidence of sulfonamide and tetracycline resistance in modern gonorrhoea is due to historic treatment of the disease itself followed by sustained use of these drugs for co-infections. The high proportion of diverse circulating strains carrying the *folP* and *rspJ* mutations could be used as evidence that they were in the gonococcal population long before the introduction of antimicrobials. However, this seems unlikely. More plausibly, the use of sulfonamides and tetracyclines has produced a strong selective pressure over an extended period of time, which has led to many independent acquisitions of resistance mutations, or more likely, convergent gains of resistance via homologous recombination. In the more recombinogenic lineage A, this has resulted in these mutations sweeping through the entire clade. Furthermore, other AMR determinants that have entered the gonococcal population more recently appear to be undergoing the same process, particularly in lineage A. The DNA gyrase A S91F substitution, which provides resistance to ciprofloxacin, is one of many resistance mutations that show extremely high levels of homoplasy in lineage A, consistent with rapid dissemination via recombination. The mosaic *penA* alleles, which reduce susceptibility to ESCs are another example. These elements were first described in *N. gonorrhoeae* around the turn of the century, but have already been independently acquired by a number of A sublineages, clearly showing that these mutations are transferring *en masse* via recombination rather than by repeated *de novo* mutation. Lineage B, on the other hand, has remained susceptible to most antimicrobials. More generally, levels of homoplasy and SNP clustering were found to be significantly higher in clade A, supporting the hypothesis that higher rates of recombination in this lineage have facilitated its high levels of AMR.

The rise of AMR gonorrhoea is generally assumed to have been facilitated by particular demographics who partake in high-risk sexual behaviours, particularly unprotected sex with multiple partners. These groups are also more often treated with antimicrobials than the general population due to frequent infection. Concordantly, we found that lineage A is associated with infection in MSM, one of the predominant risk groups, while isolates from B are more rarely found in this demographic group. Thus, lineage A isolates have the means (increased homologous recombination), motive (higher antimicrobial exposure) and opportunity (higher rates of coinfection with commensal *Neisseria* and other STIs) for recombination-driven gain of AMR.

Most recent media attention and gonococcal genomics research has focussed on the increasing levels of AMR in gonorrhoea. However, we have shown that a mostly susceptible lineage of gonorrhoea is successfully persisting in lower-risk groups where it is less likely to be exposed to antimicrobials. Notably, this lineage was associated with heterosexual groups, and therefore with infections in women, where rates of asymptomatic infection are high. Turner *et al*.^32^ showed, using a modelling approach, that in a situation where both resistant and susceptible strains are present in a population, high rates of asymptomatic infection, and therefore under-treatment, can allow susceptible isolates to survive and thrive. In such circumstances, rates of susceptible infection can be hugely underestimated, potentially meaning that our understanding of gonococcal prevalence, and rates of AMR may be biased. Interestingly, the majority of our African samples were from lineage B, consistent with epidemiological studies that describe a hidden epidemic of gonococcus in rural South African women, in which 48% of cases were asymptomatic and another 50% were symptomatic but not seeking care^33^. Similarly in Namibia, prevalence of asymptomatic gonococcal infections in both men and women in rural villages are high^34^. This may suggest that lineage B is associated with asymptomatic infection more fundamentally than simply being more often found in women, reminiscent of *Chlamydia trachomatis*, another highly prevalent STI which exhibits no AMR. In such a situation, if compensatory mutations are not developed, gain of AMR determinants may be detrimental as these elements may come with an associated general cost to fitness. Grad *et al*^35^ reported, for example, that 23S mutations associated with azithromycin resistance led to reduced ESC MICs in isolates with mosaic *penA* alleles. Similarly, we have observed that the *tetM*-containing pConjugative and pBlaTEM resistance plasmids are negatively associated with isolates with mosaic *penA* alleles, again suggesting these elements are detrimental in combination.

In conclusion, in the first phylogeographic analysis of a global collection of gonococci we have shown that although the modern gonococcal population is highly mixed, this mixing is relatively recent. The modern population originated as late as the 16^th^ century, most likely in Europe or Africa, and an early single introduction into Asia led to a rapid spread throughout the continent and the rest of the world. Despite most recent focus being on gonococcal AMR, we have demonstrated that *N. gonorrhoeae* has adapted to sexual networks with different risk profiles and exposures to antimicrobial treatment. Modern global gonorrhoea can be divided into two lineages, which we term A (after the phylogenetic breakpoint) and B (before the phylogenetic breakpoint). Lineage A has gained and proliferated AMR determinants, aided by an increased rate of recombination. These isolates are often transmitted in higher-risk networks, e.g. MSM, where pharyngeal infections are more common and individuals are often exposed to treatment for gonorrhoea and other STIs. Lineage B, however, has not gained AMR so rapidly, with 26% of isolates containing no known AMR determinants, and is potentially being silently transmitted in undertreated groups where levels of asymptomatic infection are higher. Thus, our results have shown that the effect of antimicrobial treatment on the gonococcal population has been more complex than simply initiating an inexorable progression towards AMR.

## Supplementary Information

Supplementary Information is linked to the online version of the paper at www.nature.com/nature.

## Acknowledgements

We thank Hien To and Olivier Gascuel for their help with the LSD software, and the Pathogen Informatics group at the Wellcome Sanger Institute for informatics support. We also thank Prof. Simon Szreter, Prof. Tim Bayliss-Smith and Dr. Piers Mitchell from the University of Cambridge for interesting discussions on the historical evidence of gonorrhoea infection. Japanese isolates were kindly provided by Yuko Watanabe and Toshio Kuroki, Department of Microbiology, Kanagawa Prefectural Institute of Public Health, Kanagawa, Japan. This work was funded by Wellcome grant number 098051 and the Foundation for Medical Research at Örebro University Hospital, Örebro, Sweden. JC was funded by the ERC grant no. 745258. YHG is supported by The Smith Family Foundation and NIH/NIAID grant 1R01AI132606-01.

## Author contributions

SRH, MU, SDB and JP conceived and managed the study. LSB and SRH analysed the data and drafted the manuscript. DG, MU and MO cultured isolates and extracted DNA. LSB, SRH, MU and YG interpreted the data. JC provided statistical analysis. RF advised on historical interpretation. All authors contributed to the writing of the manuscript.

## Author information

Reprints and permissions information is available at www.nature.com/reprints. The authors declare no competing interests. Readers are welcome to comment on the online version of the paper. Publisher’s note: Springer Nature remains neutral with regard to jurisdictional claims in published maps and institutional affiliations. Correspondence and requests for materials should be addressed to S.R.H.

## Figure and table legends

**Supplementary Table 1 | Additional strain information.** Metadata associated to the 419 *N. gonorrhoeae* strains included in the study, typing information and antimicrobial resistance determinants detected by ARIBA.

**Supplementary Table 2 | Analysis of Molecular Variance.** Analysis of Molecular Variance (AMOVA) calculated using three population levels: continent, subcontinent and country, for real and randomized population structure. Randomization shows no population structure at all, supporting the signal obtained by our data.

**Supplementary Table 3 | Multivariate Analysis of Variance.** Results of the Multivariate Analysis of Variance (MANOVA, Wilk’s lambda test) calculated to assess the significance of each discriminant function in the DAPC analysis. ****p-value<0.0001.

**Supplementary Table 4 | Prior and posterior continent assignments.** Prior and posterior assignments to continents for each of the strains included in the study after running a DAPC analysis.

**Supplementary Table 5 | Membership assignments by continent.** Number of prior and posterior membership assignments to each continent after DAPC analysis.

**Supplementary Table 6 | Root ancestry.** Percentage of ancestry of the root node of the tree to each of the five continents (posterior assignments). Summary of 1000 stochastic maps. The analysis was repeated by incorporating extra American strains from Grad *et al* 2014^15^. The number of strains with a posterior assignment to each continent used in each analysis is specified between brackets. Even sampling was obtained by subsampling the strains from different continents 100 times and performing 10 stochastic maps on each of them. Continents were randomised on an evenly sampled tree including extra American strains and the analysis repeated using 1000 stochastic maps. The highest percentage is highlighted.

**Supplementary Table 7 | Estimates of the substitution rate and tMRCA from BEAST and LSD.** Estimates of substitution rate and time for the Most Recent Common Ancestor (tMRCA) for five BAPS clusters obtained with BEAST and LSD. R^2^ shows the correlation coefficient between the root-to-tip distances and dates of isolation. HPD = High Posterior Density.

## Methods

### Global *N. gonorrhoeae* strains and antimicrobial susceptibility testing

A total of 413 *N. gonorrhoeae* strains without known epidemiological relatedness were collected from patients suffering gonorrhoea in 58 countries spanning five continents (Supplementary Table 1). Six genome references were also included in the study, spanning a range of isolation dates between 1960 and 2013 in total. Bacterial isolation from the corresponding samples, preservation and transportation was performed following standard microbiological procedures^44^. β-lactamase production and minimum inhibitory concentrations (MIC) were tested for a range of antimicrobials as described previously^45^: spectinomycin, tetracycline, penicillin G, ciprofloxacin, azithromycin, cefixime and ceftriaxone.

### DNA preparation and whole-genome sequencing

All isolates were confirmed to be *N. gonorrhoeae* and genomic DNA was extracted from the isolates using the Promega Wizard DNA purification kit, following the instructions from the manufacturer. Purified DNAs were multiplexed and sequenced using two lanes of the HiSeq 2500 2×100 bp platform at the Wellcome Sanger Institute.

### Mapping and variant calling

Fastq files from the 413 new gonococcal strains and the *N. meningitidis* 10356_1#65 outgroup (ENA accession ERS248641) were mapped to a common reference, *N. gonorrhoeae* FA1090 (NCBI accession NC_002946, 2,153,922 bp) using SMALT v0.7.4 (http://www.sanger.ac.uk/science/tools/smalt-0). Variants were called using SAMtools and BCFtools v1.2^46^ after indel realignment with GATK v1.5.9^47^ and further filtered as described previously^48^.

Six public reference genomes were obtained from the NCBI and aligned using progressiveMAUVE v2.3.1^49^. The XMFA output alignment was converted into a plain fasta format using *N. gonorrhoeae* FA1090 as reordering reference through a custom Perl script. Positions with gaps in this reference were removed, so that the resulting alignment had homologous positions to the 2,153,922 bp in the FA1090 genome. This alignment was added into the one resulting from mapping the 413 isolates, producing a 419-strains alignment, which was used for further analyses.

### Recombination removal and phylogenetic reconstruction

Phages described in the *N. gonorrhoeae* FA1090 strain^50^ were masked in the alignment before running Gubbins^51^, which was used to identify genome-wide recombinant segments using high single nucleotide polymorphism (SNP) density as a marker. The *N. meningitidis* 10356_1#65 strain was used as outgroup so that events affecting all *N. gonorrhoeae* strains were not excluded from subsequent calculations.

The detected recombination events and repeat regions inferred by repeat-match (MUMMER v3.23)^52^ on the *N. gonorrhoeae* FA1090 strain genome were masked in order to minimize the occurrence of false-positive SNPs. Gblocks v0.91b^53^ was run on the resulting alignment to further clean poorly aligned regions that may introduce noise to phylogenetic analyses. Gblocks was run by allowing gap positions in up to 50% of the sequences, with a minimum block length of 10 and 8 as maximum number of contiguous non-conserved positions. The resulting 1,211,180 bp clean alignment included 15,562 variable sites, identified by snp_sites^54^, and was used for population structure analysis, phylogenetic inference and divergence estimation. Genetic clusters were obtained from the non-recombining alignment using hierBAPS^55^.

The final SNP alignment was used for Maximum Likelihood (ML) phylogenetic tree reconstruction using RAxML v7.8.6^56^under the GTRGAMMA model of nucleotide substitution and 100 bootstrap replicates. Ancestral states of all SNPs before recombination removal were reconstructed onto the resulting phylogenetic tree using ACCTRAN transformation in python. Homoplasic sites in the terminal branches of the tree were detected and evaluated for the two main lineages.

### Genome de novo assembly and *in silico* typing

In parallel to the mapping process, reads were assembled using the assembly and improvement iterative pipeline developed at the Wellcome Sanger Institute^57^. Multi-locus sequence typing (MLST)^58^ and *N. gonorrhoeae* multi-antigen sequence typing (NG-MAST)^59^ typing schemes were retrieved directly from the sequences using the get_sequence_type script (https://github.com/sanger-pathogens/mlst_check/blob/master/bin/get_sequence_type) and NGMASTER^60^, respectively. The presence of the β-lactamase (*bla*^TEM^) and tetracycline (*tetM*) genes on plasmids and the Gonococcal Genomic Island (GGI) were detected using BLAST v2.3.0+^61^and ARIBA v2.4^62^. Typing was performed for the conjugative plasmid and the *bla*^TEM^ plasmids using an *in-silico* PCR (https://github.com/simonrharris/in_silico_pcr). Primers to differentiate between the Dutch and the American *tetM*-containing plasmids were obtained from Turner *et al*, 1999^63^. To type the *bla*^TEM^ plasmids, primers described in Dillon *et al*, 1999^64^were used and the resulting amplicon sizes evaluated to differentiate among the Asia, Africa and Toronto/Rio types (Supplementary Table 1).

### Analysis of population structure

In order to study population structure from the resulting alignment, the *poppr* R package^65^ was used to perform an AMOVA test on the non-recombining section of the genome^66^ on three geographical hierarchies: continent, subcontinent and country, to calculate the percentage of observed variance within and between groups. In order to test if the observed differentiation between continents was significant, a randomization test (N = 1,000 permutations) was performed using the *randtest* function from the *ade4* R package^67^, which randomly permutes the population structure to assess the observed signal of differentiation.

To further study population structure, a Discriminant Analysis of Principal Components (DAPC, *adegenet* R package)^36,68^analysis was applied to the non-recombining 15,562 SNPs alignment using continent of isolation as population. The procedure followed by this multivariate discriminant analysis tries to maximize the discrimination between the predefined groups. To avoid over-fitting and keep enough discrimination power, the optimal number of principal components (PC) to retain was determined using the a-score optimization test, which uses randomized groups to calculate the proportion of successful reassignments corrected by the number of retained PCs. This methodology resulted in 83 principal components as optimal to keep a balance between discrimination power and over-fitting. Prior assignment to continents was randomized and the DAPC analysis repeated to confirm that the observed separation among clusters does not occur by chance. Four discriminant functions were kept for the analysis, considering the number of variables was 5 continents. A Multivariate Analysis of Variance (MANOVA) test^69^ was applied to test whether there were differences between the means of the different clusters (continents) on the discriminant clustering. Wilks’ lambda was used to test the significance of this MANOVA test. Resulting p-values were adjusted for multiple tests using False Discovery Rate (FDR)^70^.

DAPC derives group membership probabilities from the retained discriminant functions. These results were used to evaluate the level of admixture in the dataset under study. Isolates assigned with >80% posterior probability to a continent different from the prior assignment were interpreted as intercontinental transmission cases. Isolates with <80% of posterior assignment to any of the continents were considered as admixed.

### Divergence estimation with LSD and BEAST

Year of isolation for all the strains was used to calculate a root-to-tip distance regression versus time to make an estimate of the temporal signal in the data. To do this, a “clustered permutation” approach was used as described^37,71^, which takes into account potential confounding temporal and genetic structure in the data. A total of 1,000 permutations were performed with this method by randomizing the isolation dates in order to get an estimate of the statistical significance of the results. This procedure was applied to the whole dataset and to the different BAPS clusters.

In order to get an estimate of the substitution rate and the Most Recent Common Ancestor (tMRCA) for the whole *N. gonorrhoeae* global collection, the Least-Square Dating (LSD) v0.3 software^38^ was used. This new approach has been shown to be robust to uncorrelated changes of the molecular clock and to give similar results to BEAST^38^. In order to compare the performance between LSD and BEAST, individual BAPS clusters were used. Specifically, Bayesian approximation using BEAST v1.8.2^39^ was run to estimate the tMRCA and the substitution rate of the genetic clusters determined by hierBAPS^55^. Three chains were run per cluster up to 100 million generations by using a GTRGAMMA model of nucleotide substitution with 4 categories, strict molecular clock with a diffuse gamma distribution (shape 0.001 and scale 1,000) and a constant population size as priors. The same configuration was used to run two different chains with the whole collection, which did not reach proper convergence because of the complexity of the dataset. LSD was also run for the BAPS clusters that reached convergence in BEAST and the results compared (Supplementary Note 1). The obtained tMRCA was further confirmed using the Wald statistic (Supplementary Note 2).

### Phylogeography with stochastic character mapping

The continent of isolation was used as a discrete trait to study changes in the distribution over the phylogenetic tree using treeBreaker^41^. This program calculates the per-branch posterior probability of having a change in the distribution of a discrete character.

Stochastic character mapping^72^ with a symmetric transition model (SYM) was applied to the phylogenetic tree to get posterior probabilities for each continent at every node using the *make.simmap* function implemented in the *phytools* R package^40^. Given a phylogeny and a set of tip states (“continent” in this study), this method uses an MCMC approach to sample character histories from their posterior probability distribution consistent with those states given a model of evolution for the mapped character^73^. This procedure was applied to the prior and posterior continent assignments excluding the admixed individuals to reduce noise from the prior metadata.

An extra set of 236 isolates from the US^15^was added to the global collection and the phylogeography analyses repeated to confirm our results. To avoid biases due to different number of strains from different continents, the combined datasets were down sampled 100 times to N=41 (the maximum number of strains with a posterior assignment to the continent with the least number of strains, Africa), except for Oceania, from which there is not more data in the public databases to include, generating 100 subtrees. Ten stochastic maps were inferred for each of those subtrees and posteriorly combined using *phytools*^40^, resulting in a total of 1000 evaluated maps.

### Evolution of antimicrobial resistance determinants

Mutations conferring antimicrobial resistance in known genetic determinants (*16S* rRNA, *23S* rRNA, *rpoB, rpsJ, mtrR, folP, gyrA, parC, parE, penA, ponA* and *porB*) as well as the presence of the β-lactamase (*bla*^TEM^) and *tetM* genes^42^were obtained for the 413 strains sequenced in this study using ARIBA v2.4^62^(Supplementary Table 1) with a custom database created for *N. gonorrhoeae* (precomputed version available in https://github.com/martinghunt/ariba-publication/tree/master/N_gonorrhoeae/Ref). Subsequent analyses were performed using R^74^: the occurrence of different antimicrobial resistance determinants before and after the change point detected by treeBreaker on the distribution of continents and the distribution of MIC values for penicillin G, tetracycline, ciprofloxacin, ceftriaxone, cefixime and azithromycin against different combinations of the genetic determinants. The average number of changes from a susceptible to a resistant state was inferred for each of the resistant determinants under study in both lineages A and B independently using stochastic mapping (100 simulations) with the *make.simmap* function implemented in the *phytools* R package^40^. The inferred number was corrected by the number of edges in each lineage: N = 586 in lineage A and N = 236 in lineage B.

As an approximation of studying the risk groups characterizing the defined A and B lineages, 263 isolates from the global collection with information on the gender of the patient were combined with 376 from two North American studies with this information available^16,25^. ARIBA v2.4 was run for the extra isolates and the obtained results joined with the ones from the global dataset. Metadata on gender and number of total resistance determinants detected per strain was plotted on a recombination-free phylogenetic tree obtained as described above and differences between the two lineages evaluated using a t-test with R^74^.

In order to confirm our hypothesis on the two lineages being associated to different risk groups and antimicrobial susceptibilities, we downloaded the phylogenetic tree of 1,054 European isolates from a 2013 Euro-GASP survey from the WGSA *N. gonorrhoeae* scheme^26^ (https://www.wgsa.net/eurogasp2013). The breakpoint between lineages A and B was detected by obtaining a combined core genome alignment of this and our global set (1,47 strains in total) using Roary v3.11.3^75^ and running a pseudo-maximum likelihood tree with the resulting SNPs^54^ with FastTree v2.1.9^76^.

### Visualization

Visualization of metadata in phylogenetic trees was performed using iTOL^77^. Mapping of the and presence/absence of antimicrobial resistance determinants detected with ARIBA were visualized using Phandango^78^.

### Data availability

All genomic data has been deposited in the European Nucleotide Archive (ENA) under project number PRJEB4024. Accession numbers for the particular strains are indicated in Supplementary Table 1.

---

**Extended Data Figure 1.**
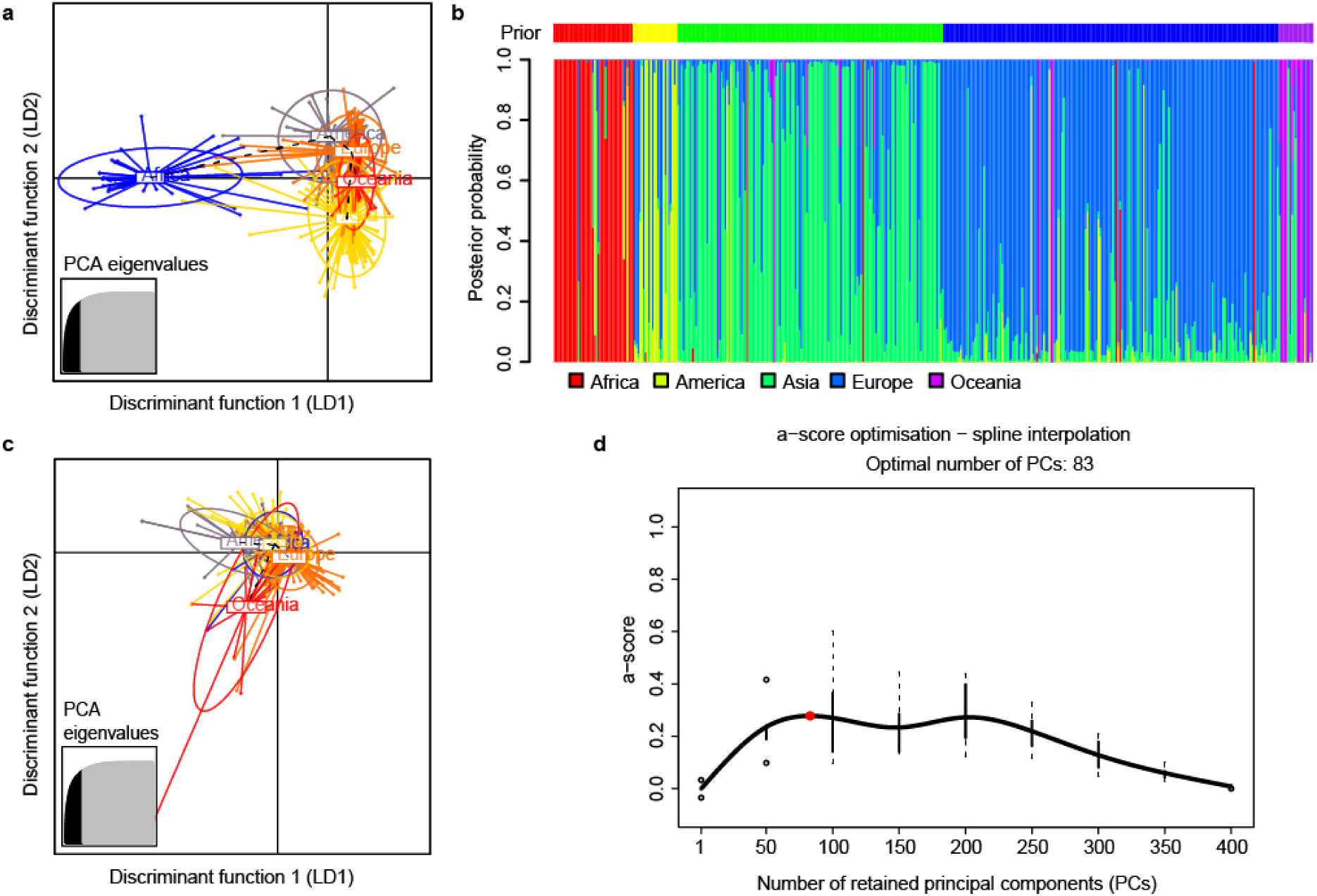
Population structure analysis. (a) Discriminant Analysis of Principal Components (DAPC) clustering of the *Neisseria gonorrhoeae* strains by its continent of isolation (p-value for discriminant function 1 < 0.001, Supplementary Table 3) (b) Membership plot showing the posterior probability assignment of each strain to each of the continents. The bar above the plot shows the prior information on the continent of isolation per strain. (c) A-score optimization test performed to obtain the optimal number of principal components to retain in the DAPC analysis^36^as a trade-off between power of discrimination and over-fitting. Calculated as the difference between the proportion of successful reassignments and values obtained using random groups corrected by the number of retained components (http://adegenet.r-forge.r-project.org/files/tutorial-dapc.pdf). (d) DAPC analysis with randomized continents showing no population differentiation by continent.

**Extended Data Figure 2.**
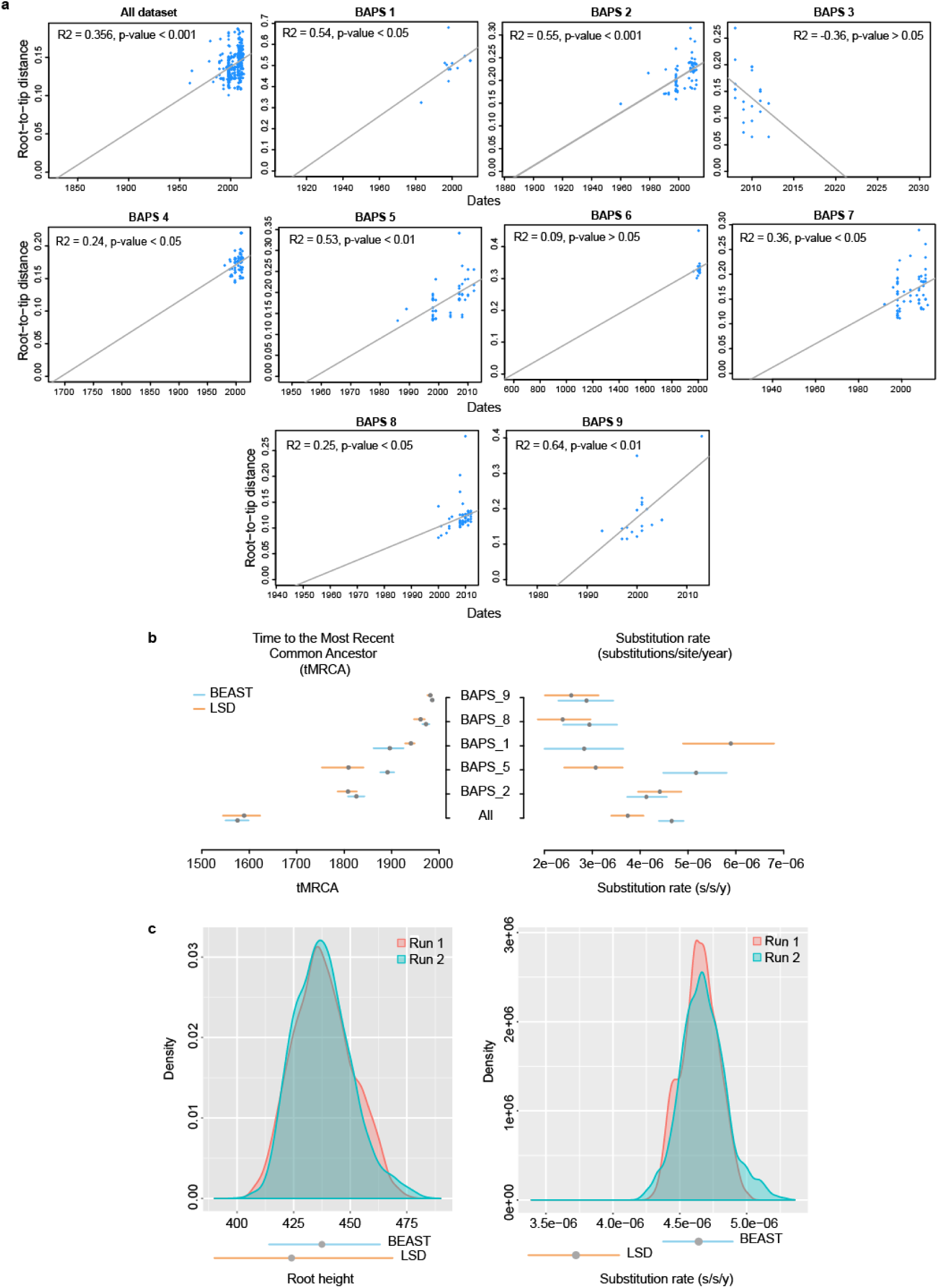
Temporal signal, tMRCA and substitution rate estimation. (a) Root-to-tip distance estimate over time of the 419 *N. gonorrhoeae* global collection (first plot) and the different BAPS clusters (following plots) calculated using the “clustered permutation” approach developed by Murray *et al*, 2017^37^. (b) Comparison between LSD^38^ and BEAST^39^. Estimates of the tMRCA and the substitution rate for the five BAPS clusters that reached convergence in BEAST as well as the whole collection calculated using BEAST and LSD. (c) Distribution of the tree root height and substitution rate parameters estimated from two different BEAST chains after 100 million generations (burn-in = 30 million). The 95% HPD interval from BEAST and the confidence interval obtained with LSD are plotted below for comparison.

**Extended Data Figure 3.**
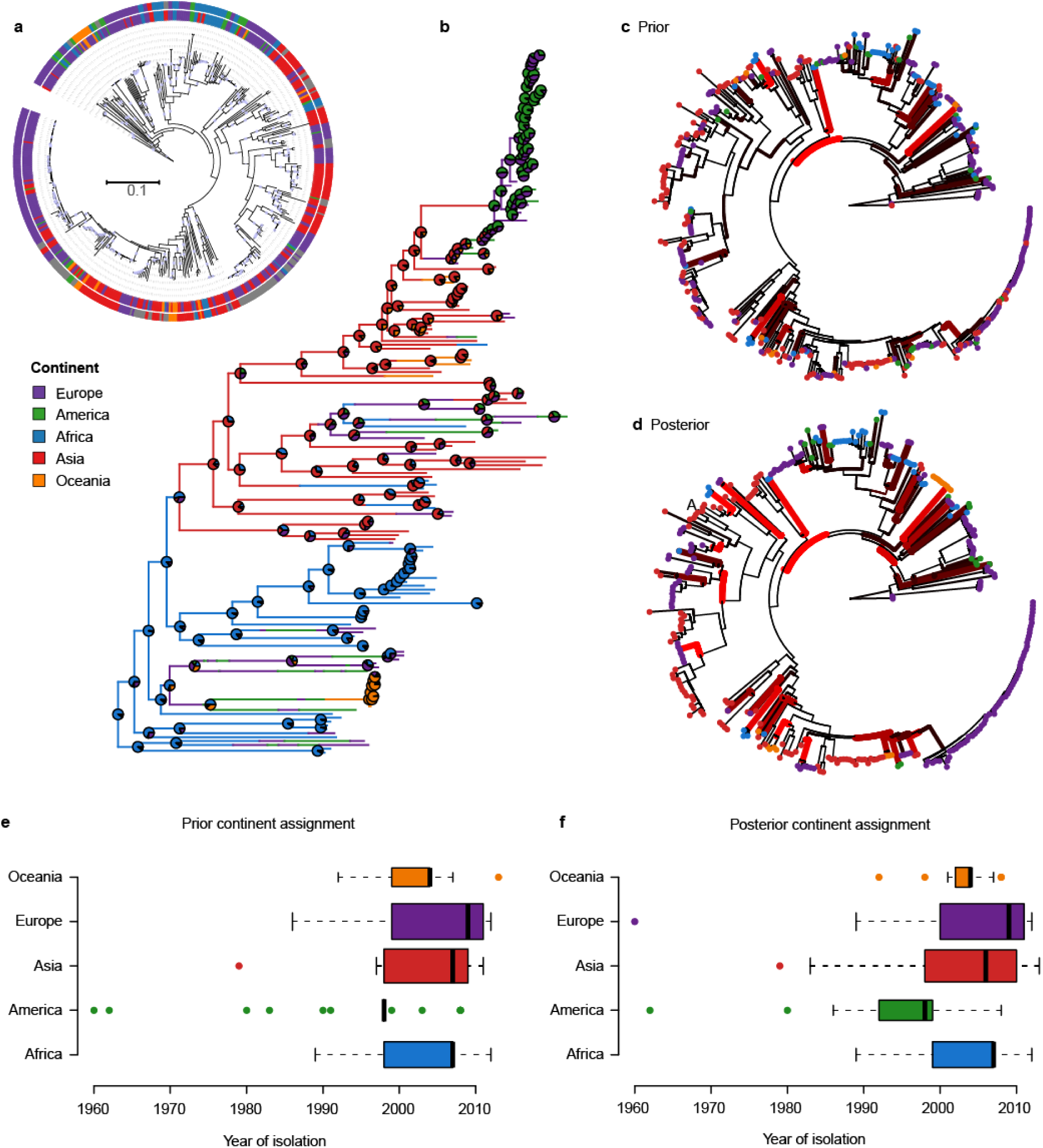
Phylogeographic analysis. (a) Maximum likelihood phylogenetic tree showing the tips coloured with the prior information on the continent of isolation (inner circle) and the posterior group membership after DAPC analysis^36^ (outer circle). Admixed individuals are defined as those with <80% posterior probability of assignment to any of the five continents and are shown in grey. Light blue circles represent bootstrap support values above 70%. (b) Stochastic mapping of the posterior continent assignments over the dated phylogenetic tree complemented with US strains from Grad *et al*. 2014^15^ and using even number of strains from each continent (N = 41) except for Oceania, for which there is not more available data in the public databases at the time of writing. Analysis was performed using the *phytools* R package^40^. Pie charts in every node represent the proportion of ancestry to each of the five continents. (c-d) Per-branch posterior distribution of the trait “Continent” using both prior (c) and posterior (d) memberships along the phylogenetic tree calculated using treeBreaker^41^. (e-f) Distribution of isolation dates by continent shown for both prior (e) and posterior (f) continent assignments. The width of the boxplots is proportional to the number of strains from each continent.

**Extended Data Figure 4.**
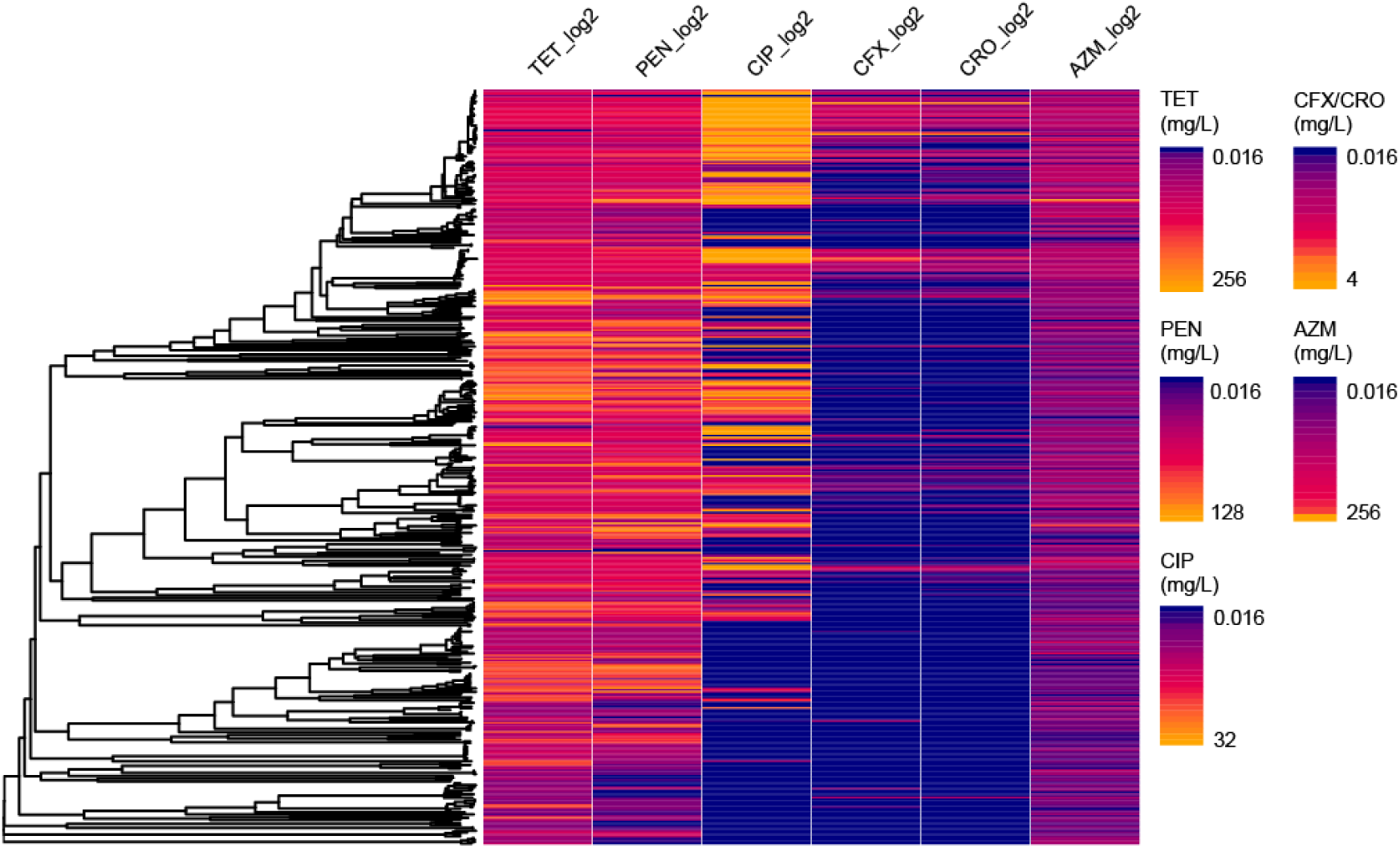
Minimum Inhibitory Concentration (MIC, in mg/L) for six antimicrobials. The distribution of the MICs of six antimicrobials is shown for all strains included in the analysis in log 2 scale. Note that the distribution of resistance is equivalent to that found from genotypic resistance. The scale of the cephalosporins CFX and CRO has been collided into one and the range of MICs in the legend is shown without logging. The minimum MIC value has been set to 0.016 mg/L for all antimicrobials. TET = Tetracycline, PEN = Penicillin G, CIP = Ciprofloxacin, CFX = Cefixime, CRO = Ceftriaxone, AZM = Azithromycin.

**Extended Data Figure 5.**
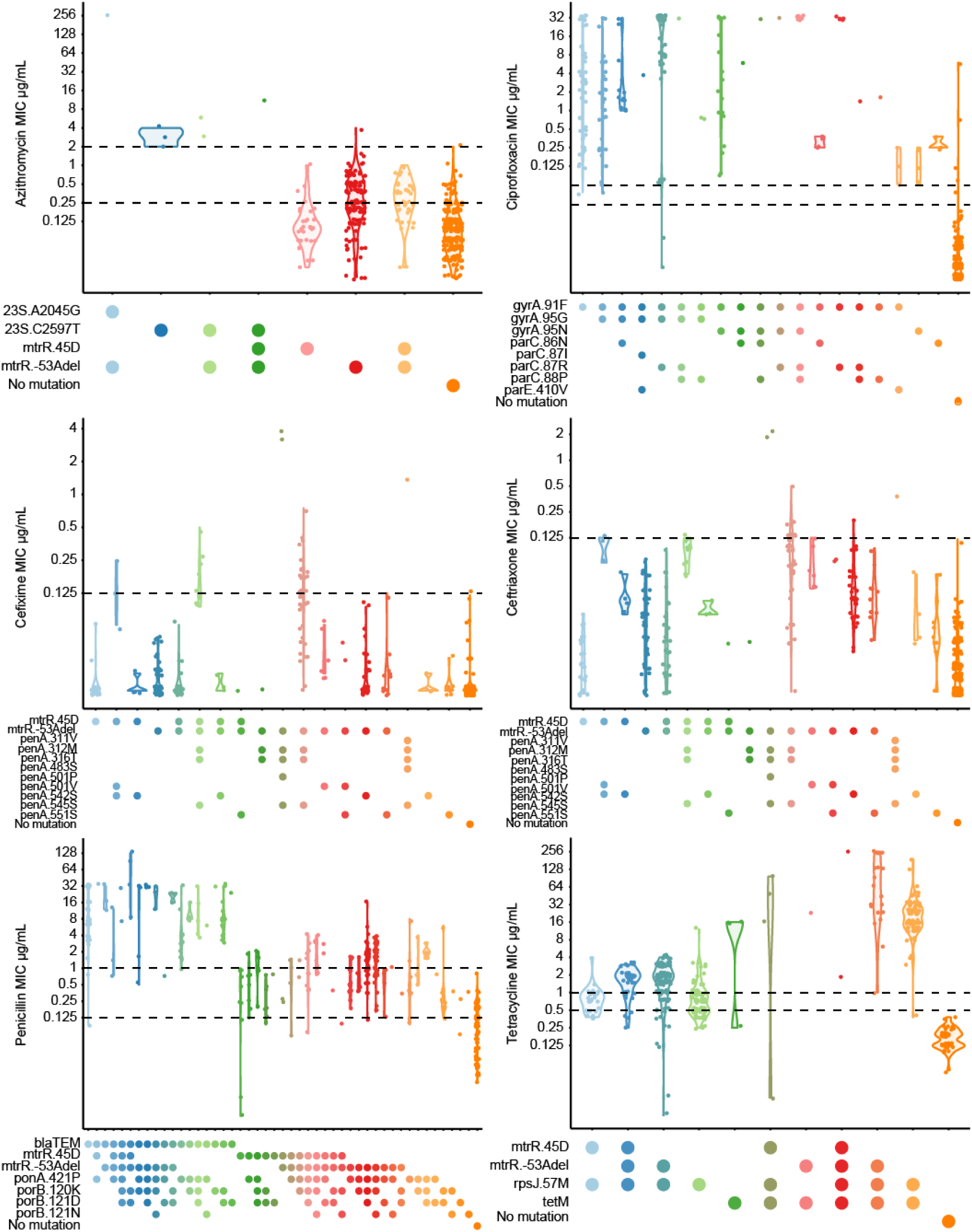
Phenotypic and genotypic resistance. Distribution of MIC values (in logarithmic scale) for each combination of antimicrobial determinants associated in the literature^42^ to each of the six antimicrobials under study. Horizontal dashed lines mark EUCAST breakpoints (www.eucast.org) except for azithromycin, in which the CLSI 2µg/mL upper bound is used (www.clsi.org).

**Extended Data Figure 6.**
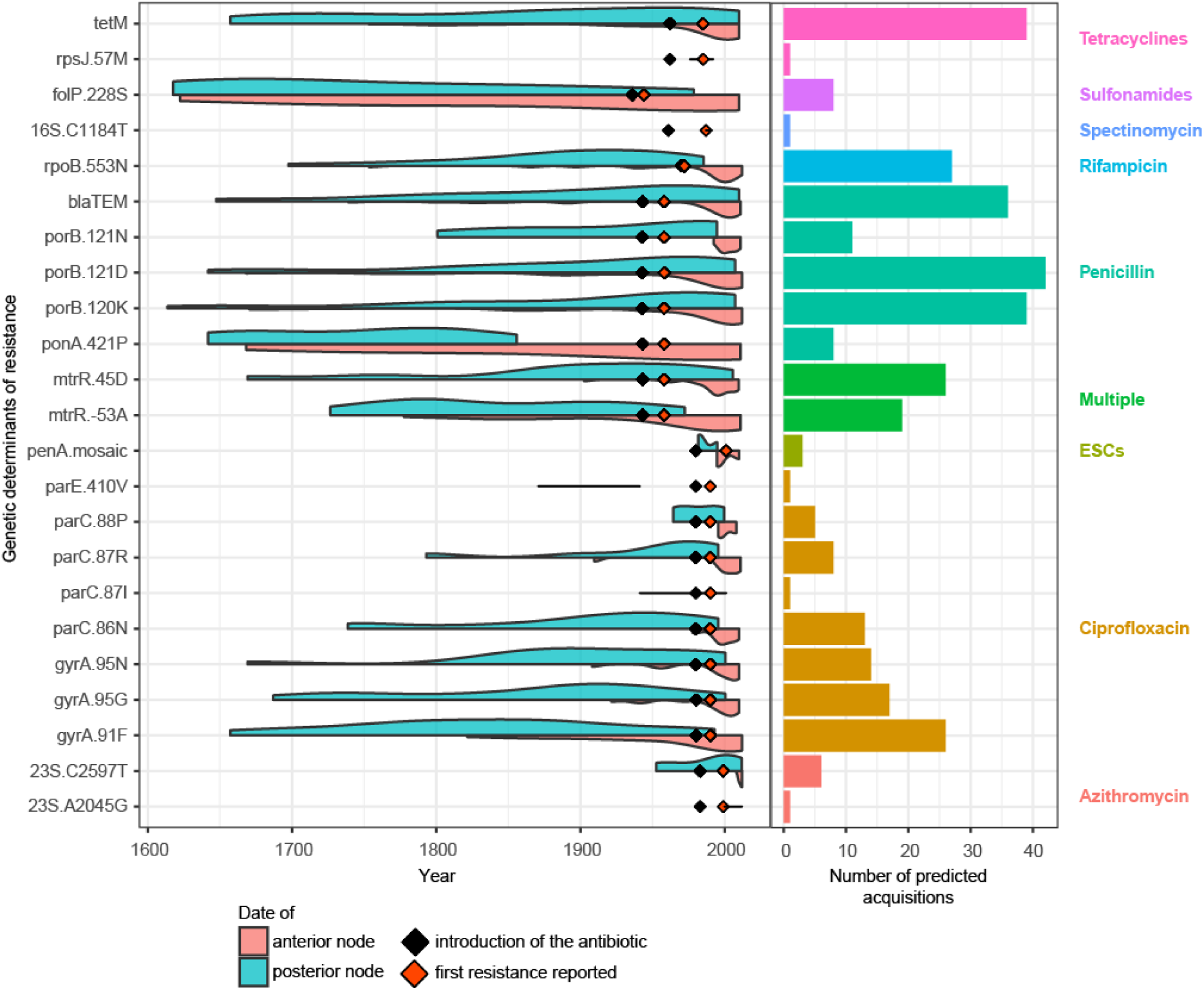
Most predicted acquisitions of antimicrobial resistance happened after the introduction of antimicrobials. The distribution of the predicted acquisitions of the known determinants of antimicrobial resistance along time are represented as half violin plots (pink when using the date of the anterior and blue the posterior nodes of the branch where the acquisition is predicted). The prediction was performed with ancestral maximum likelihood reconstruction (*ace* function of the *phytools* R package^43^) and a symmetric model of transition between states. Black diamonds mark the reported date of introduction of different antimicrobials to treat gonorrhoea and orange diamonds the first reported date of treatment failure. The barplot on the right shows the total number of predicted acquisitions for each mutation, and it is coloured to differentiate mutations affecting different antimicrobials.

**Extended Data Figure 7.**
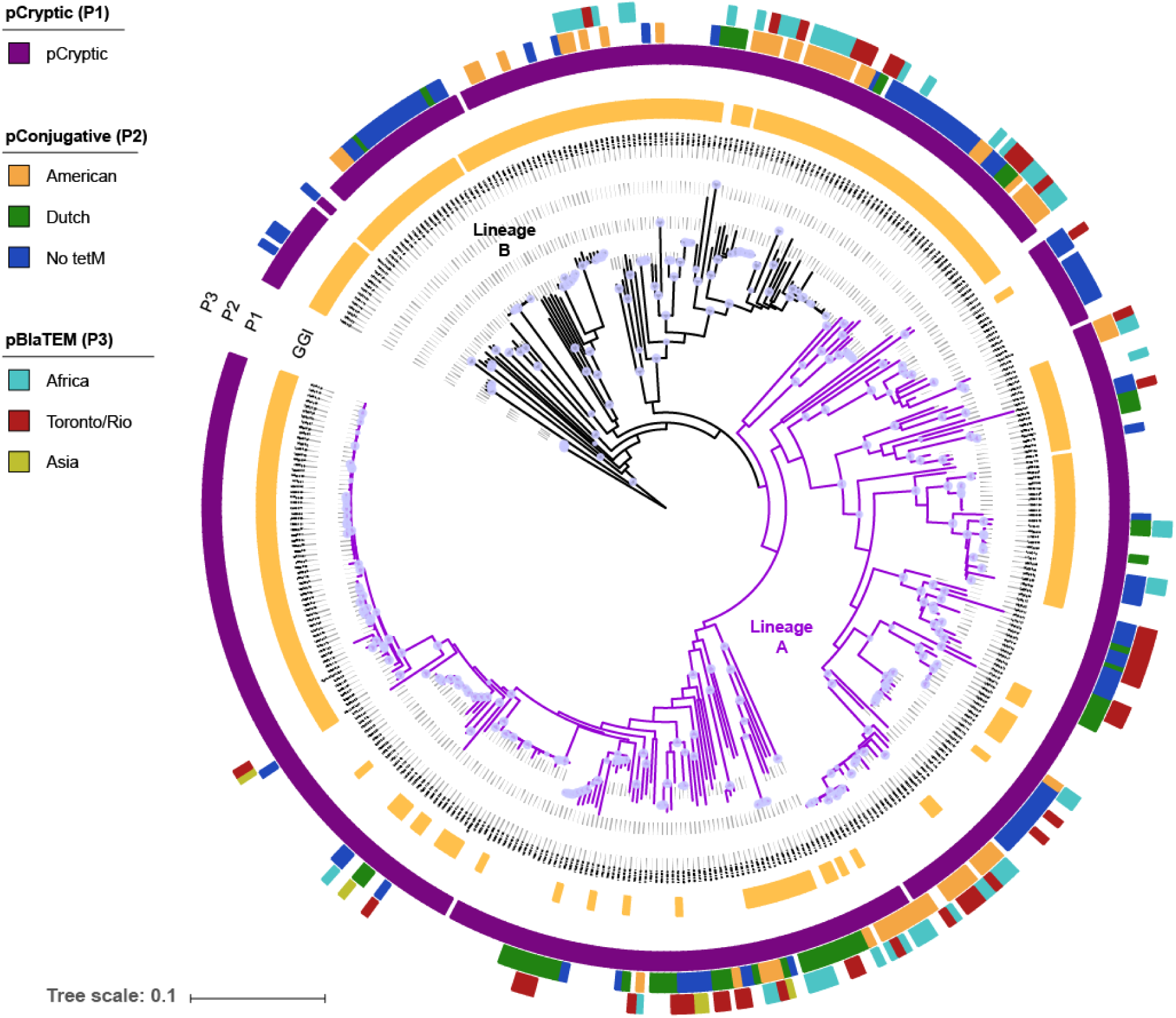
Distribution of the Gonococcal Genomic Island and plasmids. Maximum likelihood phylogenetic tree showing, from the inner to the outer circle: the occurrence of the Gonococcal Genomic Island (GGI) and the three main plasmids (pCryptic-P1, pConjugative-P2 and pBlaTEM-P3). Colours in plasmid tracks correspond to different types (see legend). The two lineages are marked in black and purple, respectively. Node shapes represent bootstrap support values above 70%.

**Extended Data Figure 8.**
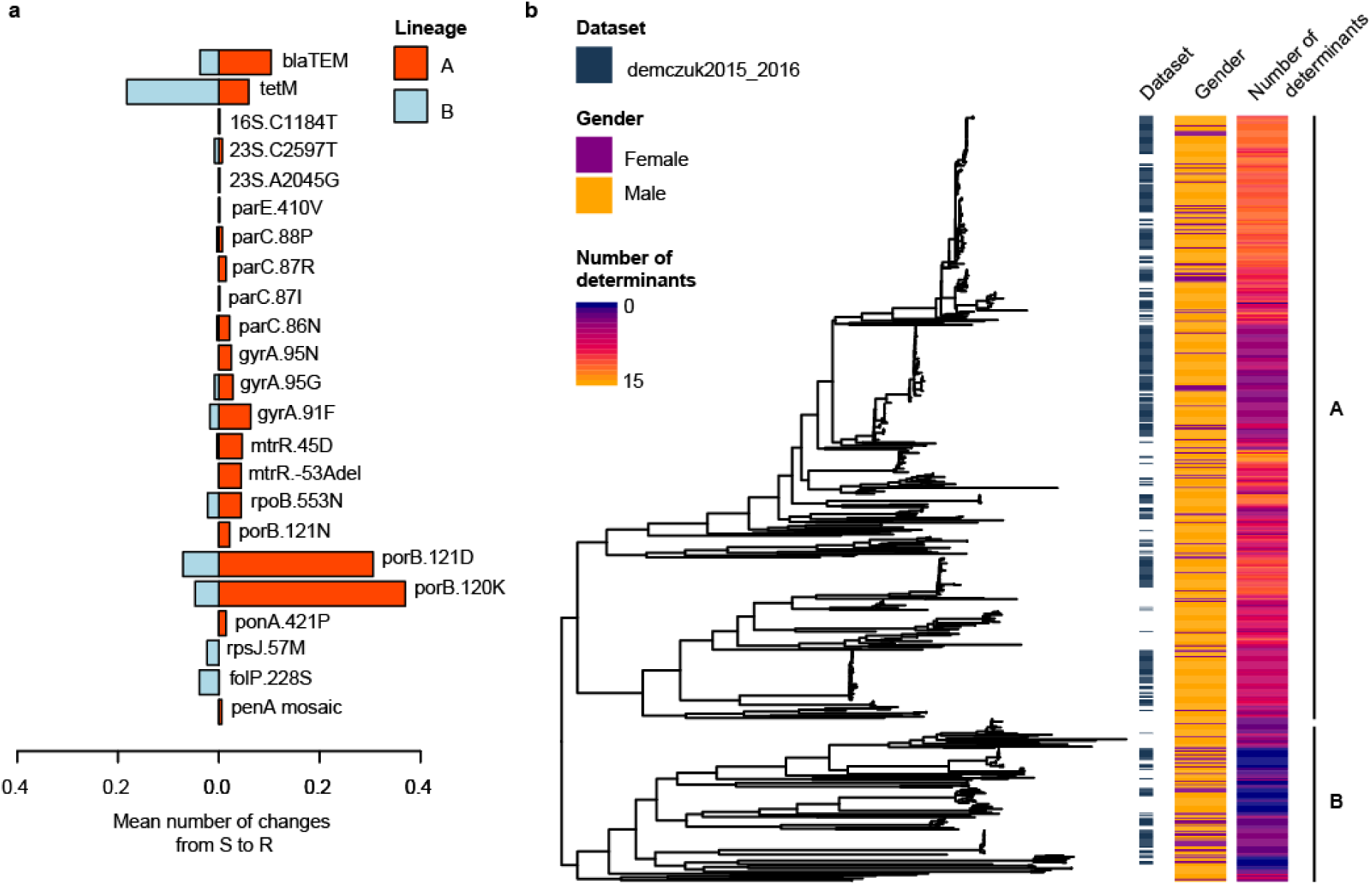
AMR genetic determinants in the two lineages. (a) Mean number of changes from susceptible to antimicrobial resistant status inferred to have evolutionarily happened in lineages A and B for all the analysed resistance determinants, including the *penA* mosaic. Values have been corrected by the number of edges of the A (N = 586) and B (N = 236) subtree lineages, respectively. (b) Distribution of the patient’s gender and the number of antimicrobial resistance determinants that the infecting strains carry. The figure shows a non-recombinant maximum likelihood tree of 639 strains (263 from the global collection and 376 from two North American studies^16,25^) with the mentioned information as metadata. Lineages A and B are also labelled.

**Extended Data Figure 9.**
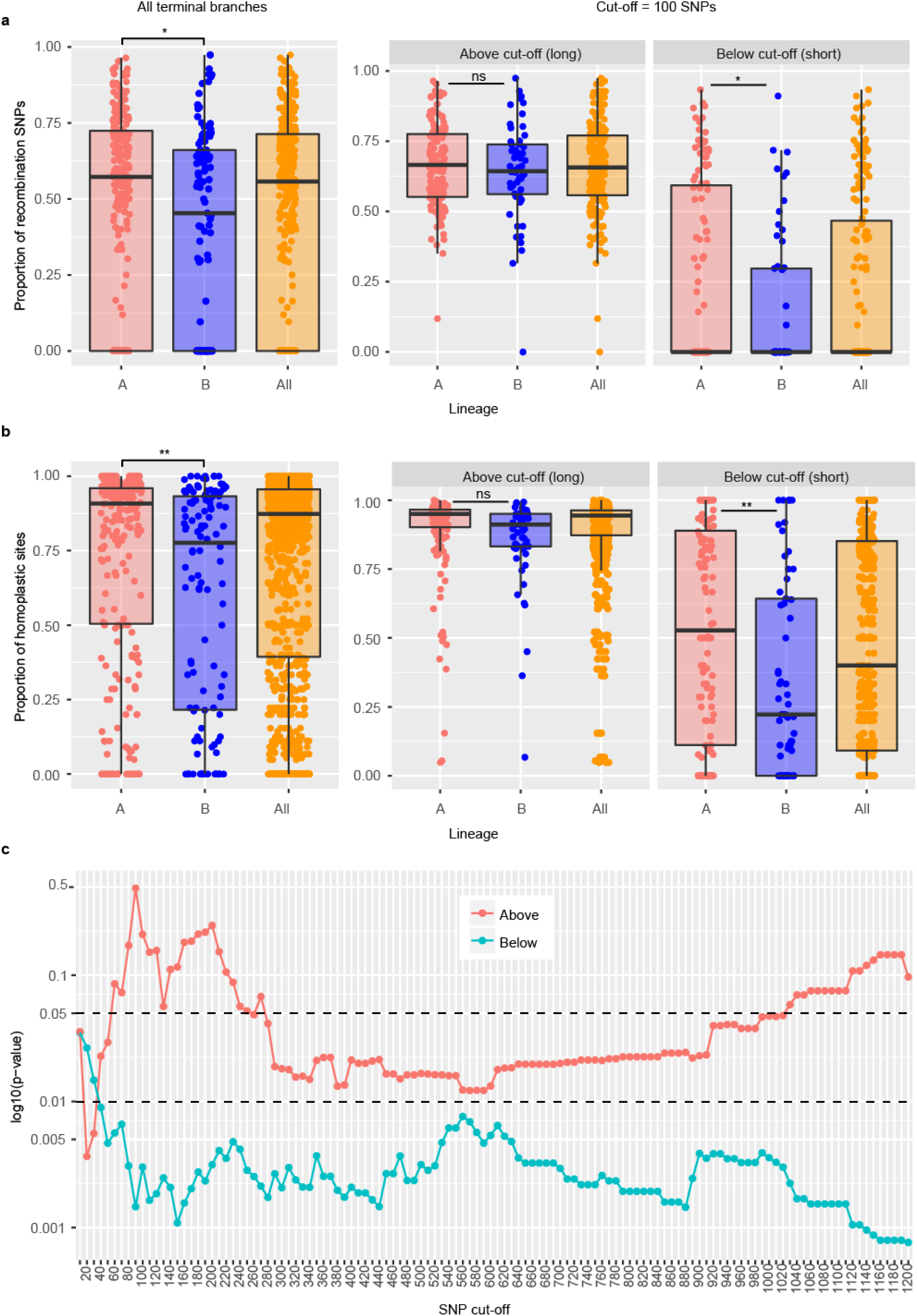
Assessment of recombination SNPs and homoplasic sites in terminal branches. (a) Proportion of SNPs inside the recombination events predicted by Gubbins for all terminal branches together and shorter (<=100 SNPs) and longer (>100 SNPs) terminal branches separately in lineages A and B and all strains. (b) Proportion of homoplasic sites in all terminal branches together and shorter (<=100 SNPs) and longer (>100 SNPs) terminal branches separately lineages A, B and all strains. (c) Distribution of the p-values calculated using a Student’s t test on the number of homoplasies in the two lineages on short and long terminal branches of the tree at different SNP cut-offs. Significance thresholds at 0.01 and 0.05 are marked with a dashed line. Short branches are probably more reliable for this type of calculation than long branches and this can be observed at around a cut-off of 100 SNPs, where branches under this number of SNPs have a p-value<0.005 while the rest are clearly not significant p-value>0.1. SNPs in repeat regions and those known to undergo antigenic variation, such as pilin-, opA54- or Maf-associated genes were excluded from the calculation. **pvalue<0.01, *p-value < 0.05, ns = non-significant.

**Extended Data Figure 10.**
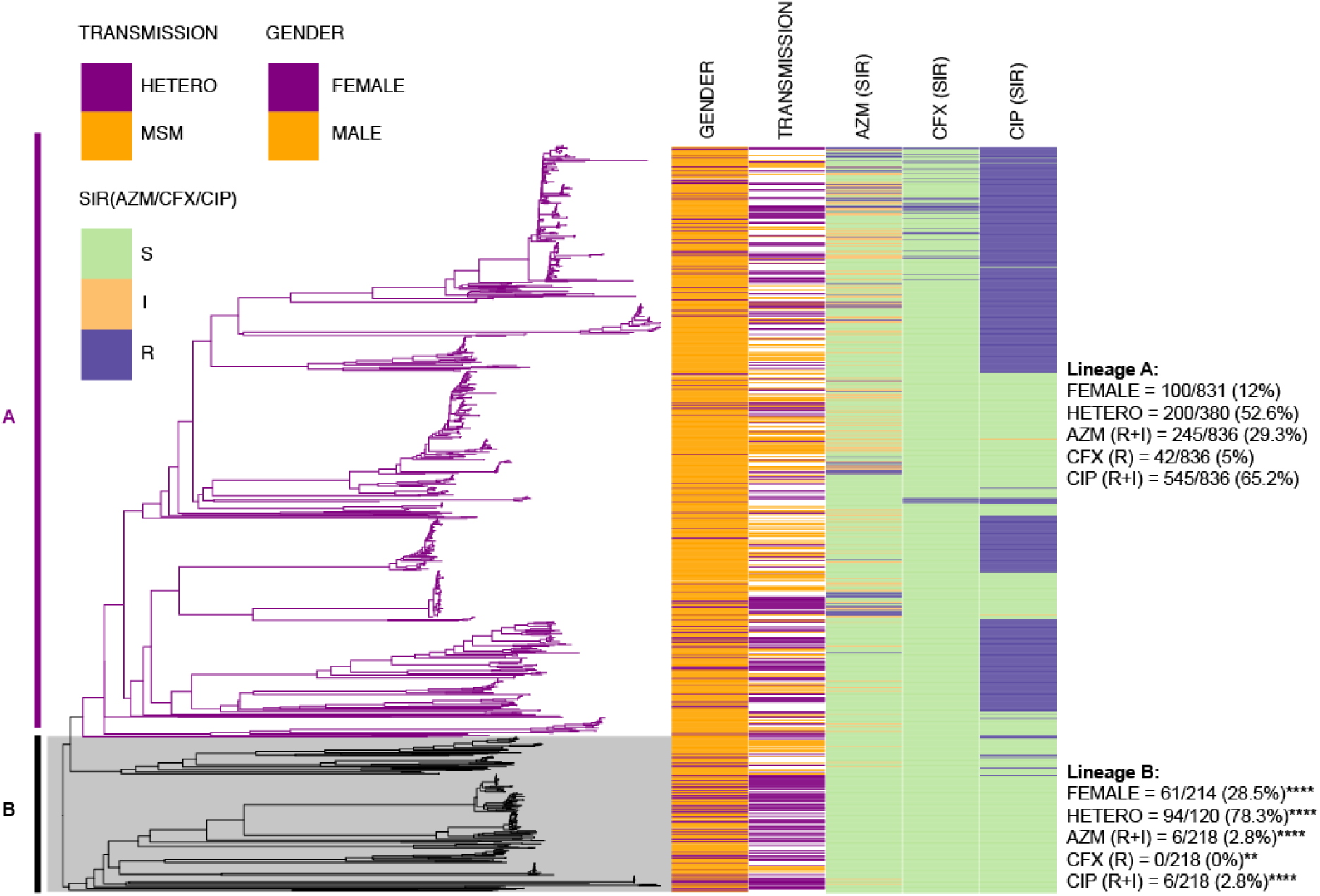
WGSA phylogenetic reconstruction of 1,054 strains from the Euro-GASP 2013 survey^26^. The two lineages (A in purple and B in black) were identified by combining this set with the global collection data as specified in the methods section. The metadata aligned to the tree shows the distribution of the gender of the patients from which the isolates were obtained, the type of transmission and the SIR categories of the phenotypic antimicrobial susceptibility test of the isolates following the breakpoints from EUCAST (www.eucast.org). Counts of each column are shown per lineage on the right side. Asterisks in lineage B indicate statistical significance compared to lineage A. AZM = Azithromycin, CFX = Cefixime, CIP = Ciprofloxacin. R = Resistant, I = Intermediate resistance, S = Susceptible. ****p-value<0.0001, **p-value<0.01.

